# Soil microbial communities in diverse agroecosystems exposed to glyphosate

**DOI:** 10.1101/484055

**Authors:** Ryan M. Kepler, Dietrich J. Epp Schmidt, Stephanie A. Yarwood, Krishna N. Reddy, Stephen O. Duke, Carl A. Bradley, Martin M. Williams, Jeffery Buyer, Michel A. Cavigelli, Jude E. Maul

## Abstract

In spite of glyphosate’s wide use in agriculture, questions remain about effects of the herbicide on soil microbial communities. Conflicting scientific literature reports divergent results; from no observable effect of glyphosate to the enrichment of common agricultural pathogens such as *Fusarium*. We conducted a comprehensive field-based study to compare treatments that did and did not receive foliar application of glyphosate spray. The study included two field sites, Maryland and Mississippi; two crops, soybean and corn; four site years, 2013 and 2014; and a variety of organic and conventional farming systems. Using amplicon sequencing, the prokaryotic (16S rRNA) and fungal (ITS) communities were described along with chemical and physical properties of the soil. Sections of corn and soy roots were plated to screen for the presence of plant pathogens. Geography, farming system, and seasonal progression were significant factors determining composition of fungal and bacterial communities. Plots treated with or without glyphosate did not differ in overall microbial community composition after controlling for these factors. No differential effect of glyphosate treatment was found in the relative abundance of organisms such as *Fusarium* spp. or putative growth-promoting bacteria *Pseudomonas* spp.

## Introduction

Providing food for the exponentially growing global human population [1] requires agricultural productivity to double by the year 2050 [2]. Thirty six percent of the Earth’s potential agricultural land is already under production, and much of the remaining land is considered marginal and susceptible to degradation when put under intensive management [3]. External inputs for nutritional supplementation and pest control are significant production costs and non-point sources of pollution that negatively impact human and environmental health. Thus, to continue feeding the world population, farmers need new approaches to increase agricultural productivity while simultaneously mitigating negative environmental impacts [2, 4].

Introduction of genetically modified glyphosate-resistant (GR) crops has transformed agroecosystems across the globe by increasing adoption of no-till agriculture where weeds are controlled chemically rather than by tillage [5]. Glyphosate interrupts the shikimate biosynthesis pathway [6], which is responsible for the production of aromatic amino acids and other key components of cell metabolism. The shikimate pathway is found in bacteria, fungi, algae, plants and some protozoans, although not in animals. Glyphosate competitively binds to the enzyme 5-enolpyruvylshikimate 3-phosphate synthase (EPSPS) and is known to be lethal to most species of plants and a large proportion fungi. However, some microbes are resistant to glyphosate due to rapid metabolism of glyphosate or to a GR form of EPSPS. Once this biosynthetic pathway is blocked, plants die due to metabolic disruption. Even at sub lethal application rates [7, 8] glyphosate can weaken a plant’s hypersensitive response enough that a pathogen is able to infect and kill the plant. In the absence of a pathogen the plant may have a stunted appearance for a few weeks but then recover.

Plants have been shown to exude glyphosate from their roots within 24 hrs of foliar application [9]. Glyphosate strongly binds to some soil components, making it rather immobile in most soil types [10]. Its tight binding to soil contributes to its weak phytotoxicity to plants as a soil applied herbicide. The episodic exudation of glyphosate may have indirect effects on the soil microbial community, and these changes may be important to the long term sustainability of agroecosystems, but patterns or changes in the microbial community are difficult to detect in the context of seasonality, changing crop species, and geographic locations.

Concerns have been raised about increased pathogen loads and suppression of beneficial organisms associated with glyphosate use [11, 12]. There are two mechanisms by which glyphosate could enrich the soil for plant pathogens: 1) pathogens can attack glyphosate-susceptible weeds that succumb to the herbicide, the dying biomass of which then acts as refugia for subsequent crop infestation (green bridge) or 2) pathogens can gain a “foothold” in a glyphosate-resistant plant due to reduced immune response from alterations in the shikimate pathway, resulting in a non-lethal infection that allows the pathogen to propagate. A review of all GR crops by Hammerschmidt [13] determined there is no conclusive evidence that glyphosate increases the susceptibility of GR crops to disease. Another review [14] challenges this conclusion. For example, several studies have observed that GR beets and soybean have increased susceptibility to pathogens when glyphosate is applied at recommended rates [14–16]. One study found no effect of glyphosate on disease induction in GR beets until rates exceeded normal field application rates by one order of magnitude [17]. However, other studies with GR crops have found no influence of glyphosate on disease [18], as well as instances of fungicidal activity against plant pathogens, especially rusts (reviewed by Duke [19]).

Two key studies have substantiated the glyphosate-pathogen-enrichment hypothesis, finding over long study periods that glyphosate repeatedly increases the rate of colonization of crops by *Fusarium* (presumed to be a pathogenic strain), while decreasing the abundance of fluorescent *Pseudomonas* bacteria (taken as putative beneficial organisms) in the soil [11, 16]. These studies are often cited as conclusive evidence that long-term use of glyphosate increases the pathogen load and decreases the abundance of growth promoting bacteria in soils. Both studies applied culture-based methodology to quantify these microbial groups, with molecular analysis of the ribosomal internal transcribed spacer region (ITS) for fungi; however, the identification techniques employed were not sufficiently discriminatory to distinguish pathogenic and beneficial genotypes for either group. Studies using culture-free methodology to characterize microbial communities have failed to detect substantial glyphosate effects on pathogen abundance [20, 21]. The key to conclusive determination of glyphosate effect on microbial communities of GR crops is to carefully compare glyphosate sprayed and non-sprayed treatments within an agronomic context. Farming systems, soil factors, crop varieties, glyphosate legacy and application rates can all impact the behavior of glyphosate and its interaction with the crop and soil microbiome [22].

We conducted a field-scale study to observe the effects of glyphosate on the soil microbiome and plant health for corn and soybean GR varieties. Specifically, we tested the hypothesis glyphosate changes the composition of the soil microbiome when controlling for differences in soils, seasonal time points, and farming systems. Furthermore, we tested the hypothesis that *Fusarium* spp. sequence abundance or culturable numbers would increase due to glyphosate treatment. Our study includes six farming systems and a total of 12 site years, representing agricultural practices as implemented on working farms. Our study targeted both naïve soil microbiomes that have not been exposed to glyphosate and those exposed to glyphosate annually. High throughput sequencing was used to generate bacterial and archaeal 16S rRNA profiles and fungal ITS profiles.

## Materials and Methods

### Description of field sites and experimental design

The study was conducted for two years at two United States Department of Agriculture, Agricultural Research Service (USDA-ARS) sites: the Sustainable Agricultural Systems Laboratory, Beltsville, MD and the Crop Production Systems Research Unit, Stoneville, MS.

The Beltsville site is managed as part of a USDA-Long Term Agricultural Research site typical of the mid-Atlantic region and described previously [23, 24]. We conducted the study in two conventional farming systems include one using a chisel plow for primary tillage (CT) and one under no-tillage management (NT). These two systems rely on mineral fertilizers, herbicides, and other pesticides as needed to manage a corn-rye cover crop-soybean-wheat/soybean rotation. One organic farming system is a three-year corn (*Zea maize*) -rye (*Cereale secale*) cover crop -soybean (*Glycine max*) -wheat (*Triticum aestivum*)/legume (*Vicia villosa*) rotation (Org3). The second organic farming system is a six-year crop rotation (Org6) in which alfalfa (*Medicago sativa*), a perennial crop in place for three years of the rotation, replaces the vetch present in Org3. The organic systems rely on legumes, poultry litter and K_2_SO_4_ to supply crop nutrients in accordance to soil test results and local regulations. A moldboard plow and/or a chisel plow is used for primary tillage in the organic systems. Weed control in the organic systems included use of a rotary hoe and between row cultivation after crops were planted.

In Stoneville the experiment was conducted in two adjacent fields, one with a legacy of glyphosate use, the other with no glyphosate history. The field with a history of glyphosate use had GR soybean and cotton (*Gossypium hirsutum*) grown in rotation for the last 15 years prior to the experiment. The field without glyphosate history had been maintained for weed biology studies in a cogongrass [*Imperata cylindrica* (L.) Beauv.] monoculture with no herbicides applied for 12 years prior to the experiment. Field preparation included killing the cogongrass with repeated tillage, planting non-GR soybean and non-GR corn for one season prior to the current field experiment, and flail mowing at maturity.

The experiment was conducted during both the corn and soybean phases of crop rotations at both sites. At each site the following glyphosate treatments were established: a GR cultivar with no glyphosate applied and the same GR cultivar with glyphosate applied at 0.87 kg ha^−1^ twice at 5 and 7 weeks after planting. Each plot was four rows (4.6 m) wide and 6.1 m long. Soybean cultivar USG Allen (GR) was planted at 350,000 seeds ha^−1^ and the corn cultivar DKC 65-17 RR2 (GR) was planted at 30,000 seeds ha^−1^. In Beltsville the corn or soybean plots are each a phase of the main plot rotation which is farming system (NT-18yr, CT-18yr, Org3-none or Org6-none) in this experimental design each phase of the rotation is considered a split-plot of the main plot which is cropping system. At both sites four replicates of each factor and level were established. All plots were hoed by hand periodically throughout the season to keep them weed-free.

In October of each year, corn was harvested with an Almaco small plot combine (Almaco, Nevada, IA); grain yield was estimated at 15.5% moisture from the two center rows of the 6.1 m plots. In 2013 the soybean was harvested with a Almaco small plot combine and in 2014 the soybean were hand harvested and threshed from 3.05 m of the two center rows. Dry weights were calculated at 13.5% moisture.

### Soil Baseline Characteristics

Beltsville soils are Coastal Plain silty loam Ultisols, consisting primarily of Christiana, Keyport, Matapeake and Mattapex soil map units. The soils at the Stoneville site were a silt loam typic Endoqualfs dominated by Dundee soil map units. At planting, soil samples from the top 15-cm depth were collected from each plot by combining soil from six or more cores (7.5 cm diameter and 15 cm depth) sampled in a semi-random pattern in a given plot. Samples were air-dried and sieved to 2 mm. The cores were collected on a diagonal line between the second and third crop rows, 3 m from each end of a given plot. Soil samples were analyzed by the Agricultural Analytical Services Laboratory at Pennsylvania State University for: pH, organic matter (OM), CEC, P, K, Mg, Ca, S, B, Zn, Mn, Fe, Cu, As, Al, Ba, Cd, Co, Cr, Ni, Pb, Se, and Sr. pH was determined in a 1:1 water dilution, OM was determined by mass loss on combustion and CEC was determined using the methods of Ross, D. and Q. Ketterings [25]. Mehlich 3 extractions were used to quantify Ca, Mg, and K; all other metals are expressed as total sorbed using the EPA 3050 method [26].

### Rhizosphere Soil and Root Sampling

At the V3 to V4 crop growth stage (4 to 6 weeks after planting) and one day prior to glyphosate application six plants and root-associated soil were excavated from each plot by removing soil monoliths with 15 cm radius from stem and 15 cm deep using surface sterilized sharpshooter shovels. This time point is referred to as “PRE-spray.” Soil monoliths were placed on a sieve and soil around the root ball was gently removed by shaking and passed through a 2 mm sieve; this soil was considered bulk soil. Soil adhering to roots after this procedure (considered rhizosphere soil) was brushed onto a 2 mm sieve using a camel hair brush. Roots were brushed thoroughly, yet not so the integrity of the root surface was compromised. The rhizosphere samples from the six plants were pooled and 5 g were added to a 15 ml falcon tube containing 10 ml of MoBio LifeGuard nucleic acid preservation solution. The contents of the tubes were mixed and frozen at −80 °C. Plants were placed at 4 °C until processed further.

Approximately twenty days after glyphosate was applied to the GR corn and soybean plots (at growth stage R2 to R3) the soil monolith sampling was repeated in the same plots. Samples were labeled “POST-spray.” Roots and adhering soils were collected and processed the same as for PRE-spray samples. At each site, sampling was determined by the developmental stage of the crop plants and was not constrained by Julian calendar dates.

### Identification of Endophytes from Roots

Two-centimeter sections of root were cut at random sixteen times from each of six fresh root systems for each treatment. The total wet weight of the 16 sections was recorded. Sections were surface sterilized for 2 minutes in 1.25% sodium hypochlorite, followed by three rinses in sterile distilled water. Sections were blotted dry on sterile paper towel and eight root sections were placed on a plate containing Komada’s Medium [27]. Plated roots were incubated in ambient light at room temperature until colonies emerged. Fungal mycelium and spores from emerging colonies were sampled and examined on a Nikon E60 microscope and identified to genus, or to broader morphological group, based on taxonomic features. Colonies of typical morphology were plated onto minimal media to induce sporulation for further identification. Colonies not producing spores were characterized as “non-sporulating.” Polymerase chain reaction (PCR) screens for ITS followed by cloning and sequencing were conducted on over 384 colonies of typical morphology to validate microscopic identification. The methods followed those described in (Chung et al. 2008). Sequences were quality checked and aligned using the DNAStar suite of software (DNAStar, Madison, WI, USA), and identified using the basic local alignment search tool and GenBank nucleotide data bank from the National Center for Biotechnology Information, Bethesda, MD, USA (http://www.ncbi.nlm.nih.gov/. Accessed fall, 2014).

### Illumina Sequencing Library Preparation from Rhizosphere Soils

Rhizosphere and bulk soils preserved in LifeGuard at −80 °C were thawed and 800 µl of each slurry was processed using a PowerSoil-htp 96 Well Soil DNA Isolation Kit (MoBio Laboratories Inc, Solana Beach, CA) according to the manufacturer’s recommendations. DNA was quantified and quality verified using a NanoDrop 2000 spectrophotometer (Thermo Fisher Scientific, Pittsburgh, PA). 16S metagenome sequencing was conducted according to the Illumina protocol Library Preparation Manual Part# 1504423 Rev. B, Illumina Inc., www.illumina.com. Five µl of cleaned adapter amplicon product for each sample were used for index PCR using the Nextera XT Index Kit (Part# FC-131-1002; 16S Metagenomic Sequencing Library Preparation Manual Part# 1504423 Rev. B, Illumina Inc., www.illumina.com). Index PCR products were cleaned according to the Illumina protocol (16S Metagenomic Sequencing Library Preparation Manual Part# 1504423 Rev. B, Illumina Inc., www.illumina.com), and 2 µl aliquots per sample from each 96-well PCR plate were pooled for the final Illumina library. For analysis, one-hundred µl of 10 nM solutions of each library pool were frozen and shipped on dry ice for analysis on an Illumina MiSeq system at the Center for Genome Research and Bioinformatics (CGRB), Oregon State University, Corvallis, OR. For the Beltsville site, where a total of four cropping systems were also sampled, a total of 512 samples were sequenced. For the Stoneville site 256 samples were sequenced.

### Bioinformatics and Statistical Analysis

#### Sequence filtering and trimming

Reads were returned from CGRB after initial quality control with standard Illumina workflows, including quality filtering and adapter trimming. Raw data is available via the AgData commons NUMBER. Scripts used in subsequent steps can be found at “https://github.com/rmkepler/FSP_script_repository”. Prior to joining paired ends and taxonomy assignment, forward and reverse primers were removed and sequences quality trimmed (-q 22) at the 3-prime end using Cutadapt (version 1.8.3). Reads lacking primer sequences or shorter than 75 bp before trimming were discarded.

#### Assembly and taxonomy assignment

The R package Dada2 [28] was used for paired end assembly and taxonomy assignment. The command “filterandtrim” was used to remove sequences with an expected error rate greater than two, and any sequences containing “N” values (unreadable bases). Error rates were estimated for forward and reverse reads. Filtered reads were then dereplicated with the “derepFastq” command. Dereplicated sequences were denoised with the “dada” command and then paired ends were merged. Chimeric sequences were removed with the command “removeBimeraDenovo”. Taxonomy was assigned to the chimera-free table of sequences with the dada2 implementation of the RDP classifier [29]. The UNITE database (version 7.2) [30] was used as the reference for identification of fungal ITS sequence variants, and silva (release 132) [31] was used for prokaryotes.

#### Community analysis

We transformed community count data into relative abundance, then calculated Bray-Curtis dissimilarity. Principal components analysis (PCA) was applied to the Bray-Curtis dissimilarity matrix using the ordinate function of the vegan package v. 2.4 [32] as implemented in phyloseq v. 1.22.2 [33] for both fungal and prokaryotic barcodes.

After subsetting by crop and site, richness and evenness were estimated from rarefied datasets of the raw sequence counts using vegan. DESeq2 v. 1.18.1 [34] was used to produce variance stabilized datasets [35]. Bray-Curtis dissimilarity for each sample was used for PCA. We used PERMANOVA to determine significance of main effects and interactions between the following factors: farming system, soil zone, glyphosate treatment, sampling date, and year. The farming system factor had 4 categories for Beltsville (CT, NT, Org3 and Org6) and 2 for Stoneville (NT_none, NT_15yr). All other factors had two categories at both sites: soil zone (bulk and rhizosphere); year (2013, 2014); glyphosate treatment (spray, no spray); sampling date (PRE glyphosate application, and POST glyphosate application). A repeated measures model based on the plot ID was used.

The effect of glyphosate treatment on microbial communities was tested with the Wilcoxon signed-rank test of differences between dates as implemented in the longitudinal plug-in for Qiime2 [36]. The test was applied separately for three measures of richness: observed, Shannon’s and Simpson’s.

#### Differentially abundant taxa

Tests for differentially abundant taxa in response to glyphosate treatment were conducted in DESeq2 using likelihood ratio tests after subsetting fungal and prokaryotic data by site, crop, and farming system. The test compared a full model including group, sampling date terms, as well as an interaction term, where group is defined as the combination of farming system and glyphosate treatment (e.g. Org3_spray) and sampling date corresponds to PRE-spray and POST-spray sampling events. The full model was compared to a reduced model lacking the interaction term. Thus, significant interaction terms would indicate sampling date and glyphosate application interacted to be important predictors of microbial abundance. This was tested for every fungal and bacterial taxon identified. Datasets with untransformed counts were used as the starting data, which were then variance-stabilized during testing.

## Results

From sequencing analysis a total of 68,964 unique fungal and 72,454 unique prokaryotic sequence variants were identified across all samples. Beltsville and Stoneville shared 13,964 bacterial and 5,740 fungal taxa. Stoneville featured 62,985 and 29,780 bacterial and fungal taxa, respectively. Beltsville featured 41,538 and 44,924 unique bacterial and fungal taxa, respectively. Fungal richness was higher for all Beltsville farming systems, compared with Stoneville, with the exception of the Shannon’s and Simpson’s diversity metrics for Org_3 (Figure 1A). Conversely, prokaryotic diversity was greater for Stoneville in all measures (Figure 1B).

**Figure 1.**
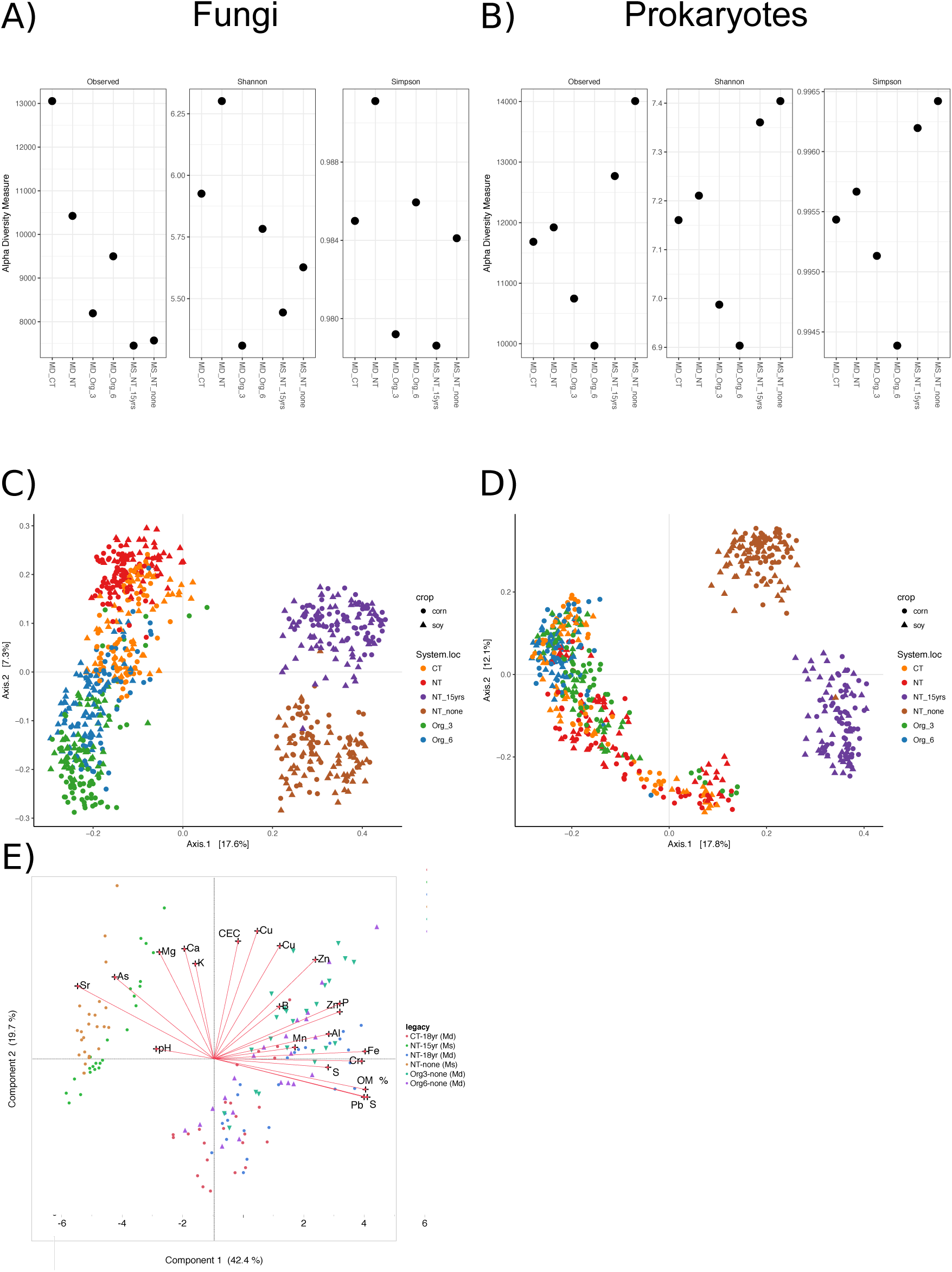
Principal component analyses of site chemistry, bacterial and fungal communities for sites in Beltsville, MD and Stoneville, MS. A) Chemical analysis of all plots in the first year of this study. B) Bray-Curtis dissimilarity of prokaryotic communities from all samples after rarefaction to a depth of 20,000 reads per sample. C) Bray-Curtis dissimilarity of fungal communities for all samples after relative abundance transformation of total counts.

Principal component analysis showed that Beltsville and Stoneville communities were distinct (Figure 1C & D). Permanova analysis of relative abundance for fungi and prokaryotes revealed that site was the most significant factor accounting for Bray-Curtis dissimilarity distances in fungi and prokaryotes (*p* = 0.001 in both cases. Fungal *R*^*2*^ = 0.19, Prokaryote *R*^*2*^ = 0.16; supplemental data). Differences between the Stoneville and Beltsville microbial communities were driven by differences in edaphic factors. Soil chemical characteristics differed between the two sites (Canonical Discrimination Analysis, p<0.001, *R*^*2*^ = 0.99), and between cropping systems (Canonical Discrimination Analysis p<0.001, *R*^*2*^ = 0.99). Soil in Stoneville was significantly higher in pH and the cations Arsenic (As), Barium (Ba) and Strontium (Anova, p<0.001), whereas Beltsville soil contained significantly more Phosphorous (P), Lead (Pb), Sulfur (S), Cromium (Cr), Iron (Fe) and OM (Anova, p<0.001) (Figure 1E). In order to increase power to detect local effects of glyphosate treatment, we analyzed sites and crop treatments separately.

Farming system was the largest driver of fungal community structure regardless of crop (Figure 2 & 3) in both Beltsville (Permanova; corn: *p* = 0.001, *R*^*2*^ = 0.16; soybean; *p* = 0.001, *R*^*2*^ = 0.16) and Stoneville (Permanova; corn: *p* = 0.001, *R*^*2*^ = 0.24; soybean; *p* = 0.001, *R*^*2*^ = 0.23). Year of sampling was also significant but explained less variance in both Beltsville (corn: *p* = 0.001, *R*^*2*^ = 0.046; soybean; *p* = 0.001, *R*^*2*^ = 0.043) and Stoneville (corn: *p* = 0.001, *R*^*2*^ = 0.051; soybean; *p* = 0.001, *R*^*2*^ = 0.052). No significant interaction was noted between sampling date and glyphosate treatment (*p* = 0.488 and 0.296 for corn and soybean, respectively).

**Figure 2.**
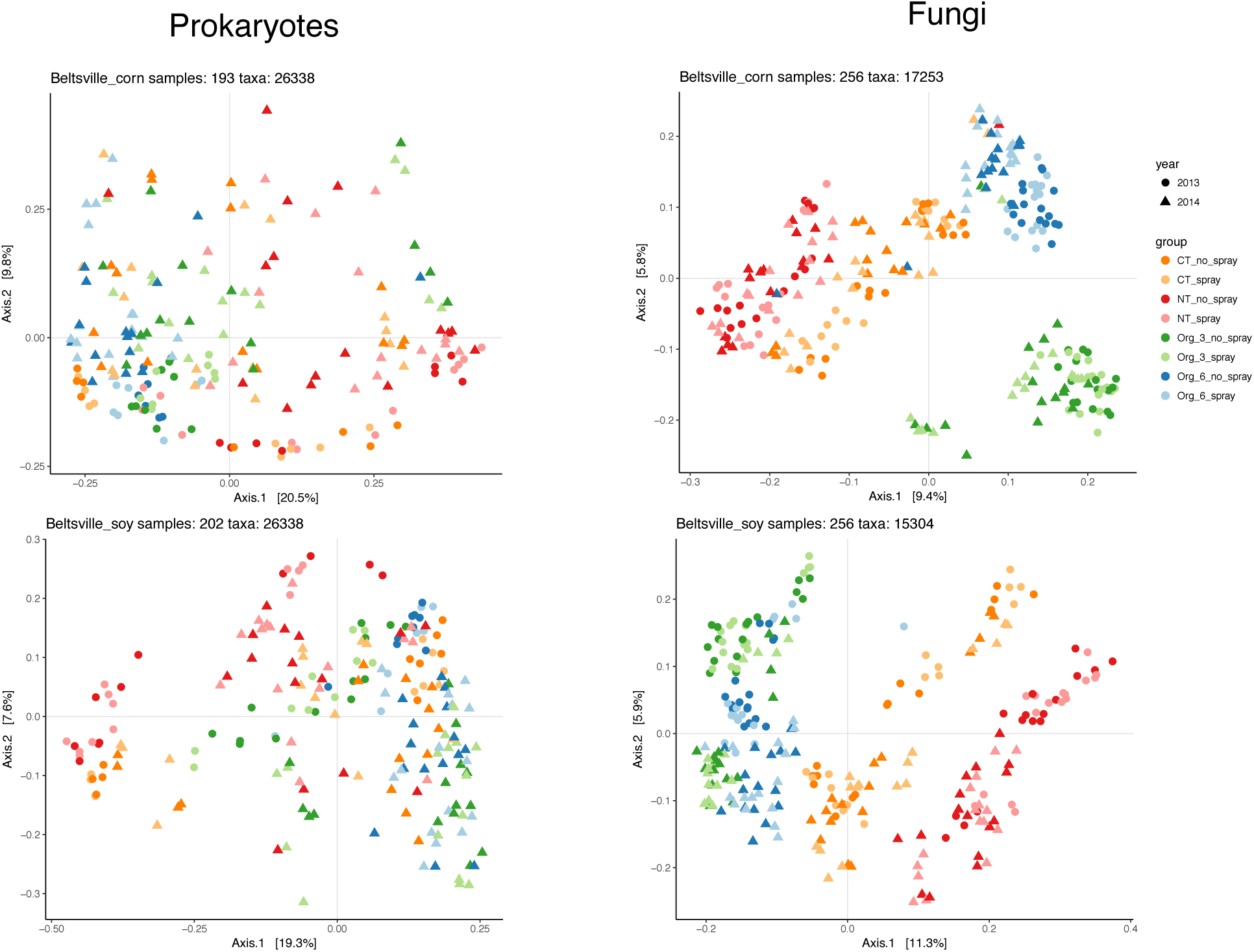
Principal component analyses of microbial communities in Beltsville, MD, partitioned by crop. Prokaryotic community data drawn from a dataset rarefied to 20,000 reads per sample. Fungal data has been variance stabilized with negative binomial transformation in DESeq2.

Rhizosphere and bulk soil samples were also not significantly different (Supplemental data) for any crop or site. Likelihood ratio tests of taxon abundance in DESeq2 also confirmed no glyphosate treatment; the sampling date-glyphosate treatment interaction did not significantly increase the explanatory power of the model for any taxon (supplementary data), regardless of crop or farming system.

Farming system was also a driver of prokaryote community structure in Beltsville (Permanova; corn: *p* = 0.001, *R*^*2*^ = 0.096; soybean; *p* = 0.001, *R*^*2*^ = 0.09) and Stoneville (Permanova; corn: *p* = 0.001, *R*^*2*^ = 0.21; soybean; *p* = 0.001, *R*^*2*^ = 0.16).Farming explained less variation in Beltsville prokaryotic communities (Figure 2), than in Stoneville (Figure 3). The year term explained a lesser amount of variance for Beltsville (corn: *p* = 0.001, *R*^*2*^ = 0.096; soybean; *p* = 0.001, *R*^*2*^ = 0.086) and Stoneville (corn: *p* = 0.001, *R*^*2*^ = 0.051; soybean; *p* = 0.001, *R*^*2*^ = 0.069). The interaction between glyphosate with sampling date was not significant for either crop (Supplemental data). Likelihood ratio tests of taxon abundance in DESeq2 also confirmed no glyphosate treatment; the sampling date-glyphosate treatment interaction did not significantly increase the explanatory power of the model for any taxon (supplementary data), regardless of crop or farming system (supplementary data).

**Figure 3.**
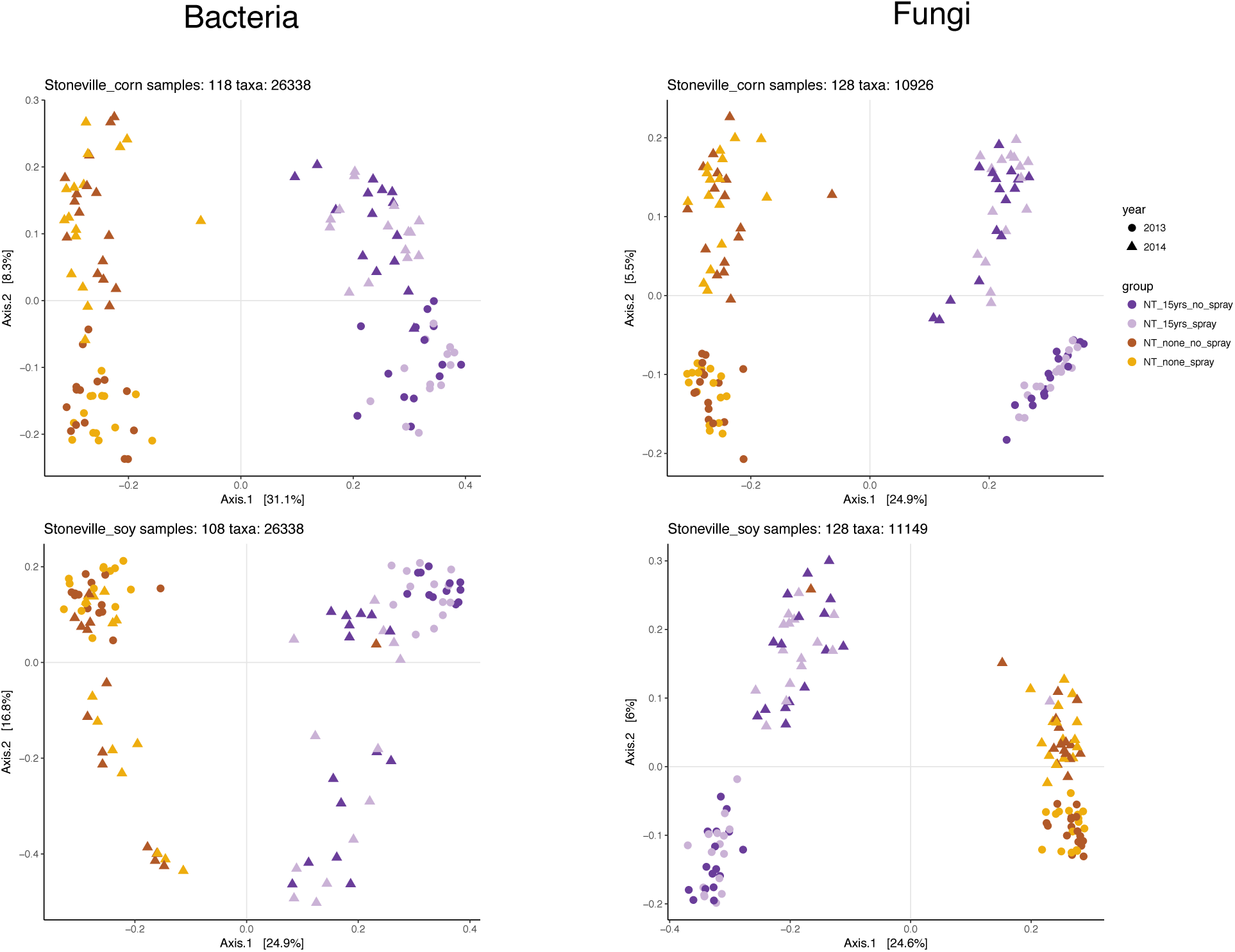
Principal component analyses of microbial communities in Stoneville, MS, partitioned by crop. Prokaryotic community data drawn from a dataset rarefied to 20,000 per sample. Fungal data has been variance stabilized with negative binomial transformation in DESeq2.

Wilcoxon rank sum tests showed several instances where species richness differed significantly between the PRE and POST sampling dates (Figure 4, Supplemental Data); however, differences were observed in both glyphosate and no-glyphosate treatments, indicating this is a seasonality effect, and not due to glyphosate exposure. In Beltsville, corn and soybean differed in their response over the two dates. Prokaryote richness for corn in every Beltsville farming system was significantly different between the two dates. This trend was also observed, but to a lesser degree in fungal communities, with half of the treatments differing significantly for both spray and no spray treatments. Fungal communities did not differ seasonally in the Beltsville soybean plots, and fungal species richness was unaffected by sampling date for both corn and soybean in the Stoneville samples.

**Figure 4.**
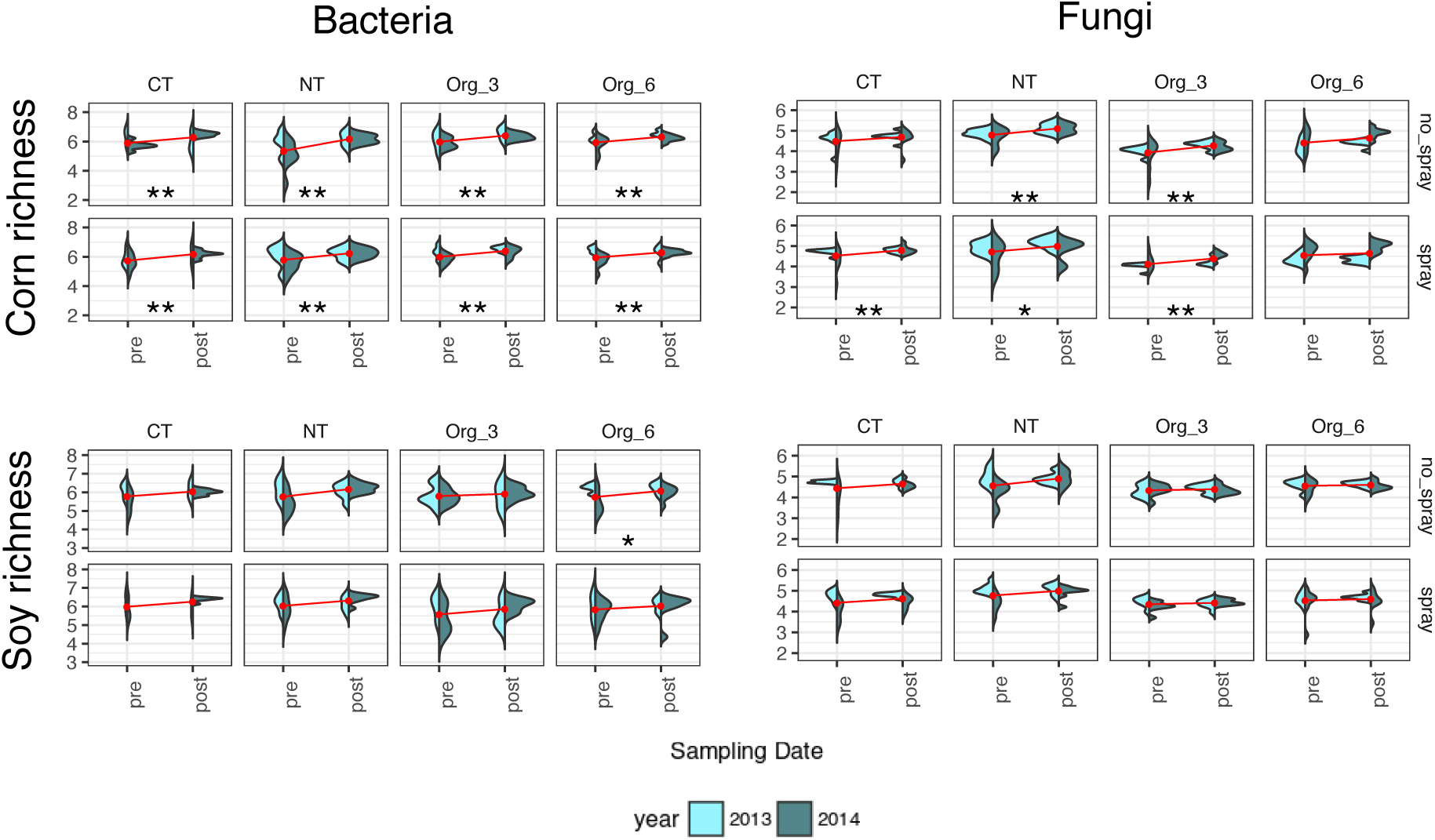
Change in Shannon’s richness of rarefied data across sampling dates in no-spray and spray treatments. Stars on each plot are for raw (*) and false discovery rate corrected (**) p-values less than 0.05 from Wilcoxon signed-rank test of differences between dates. Years are pooled, although graphed separately. Red points and line represent mean richness

**Figure 5.**
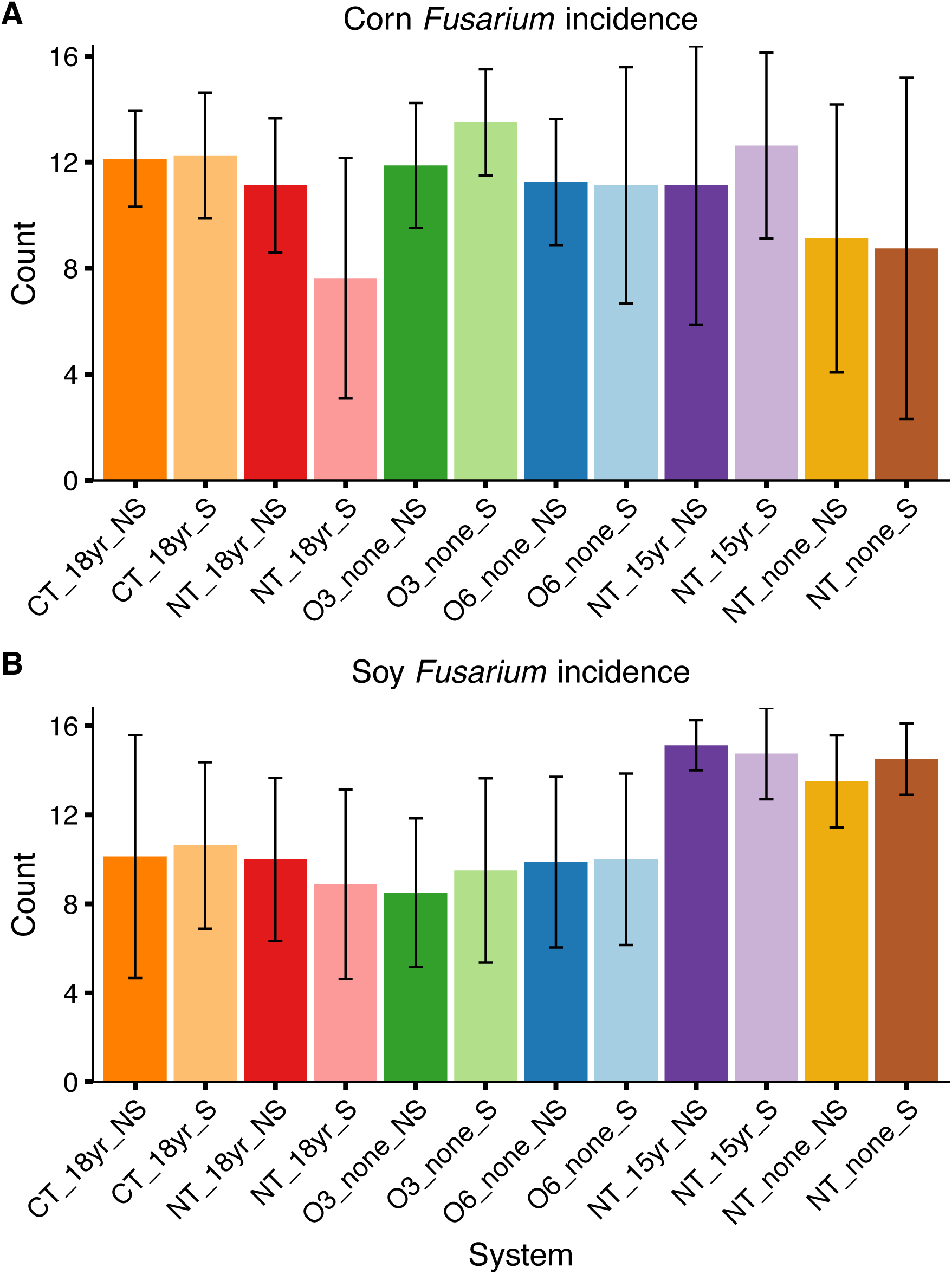
Abundance of *Fusarium* isolates +/− standard deviation. A) corn. B) soy roots. Colors follow those used in Figures 2 & 3.

The root endophyte screening required analysis of over 6100 root segments and identified over 2400 fungal colonies. Significantly more colony forming units (CFU)s were observed in 2013 than in 2014 at the Beltsville site (p<0.0003), but no differences in the number of CFUs were observed between years at the Stoneville site. A total of 384 of the typical morphotypes were ITS amplicon sequenced, resulting in 11 identified dominant taxa: *Fusarium sp., Macrophomina sp., Alternaria sp., Cladosporium sp., Penicillium sp., Zygomycota sp., Trichoderma sp*., and *Epicoccum sp*. There was no significant difference in abundance of *Fusarium sp*. or the other taxa in the glyphosate sprayed and unsprayed plots, regardless of site, crop, or year (p>0.07) (Figure 4b).

There was no significant difference in corn yield among systems or among glyphosate application treatments for either 2013 or 2014 (Table 1). Corn yields were not significantly different from the county averages for all systems with a mean among systems of 9339.4 kg/ha. In 2013 an error occurred while using the small plot combine and beans harvested from different replicates were mixed, rendering the data unusable. In 2014 soybean yields were similar to the county averages with a mean of 2326.5 kg/ha. There was no significant difference in yield across farming systems, and no effect of glyphosate treatment on yield (Table 1)

**Table 1.**
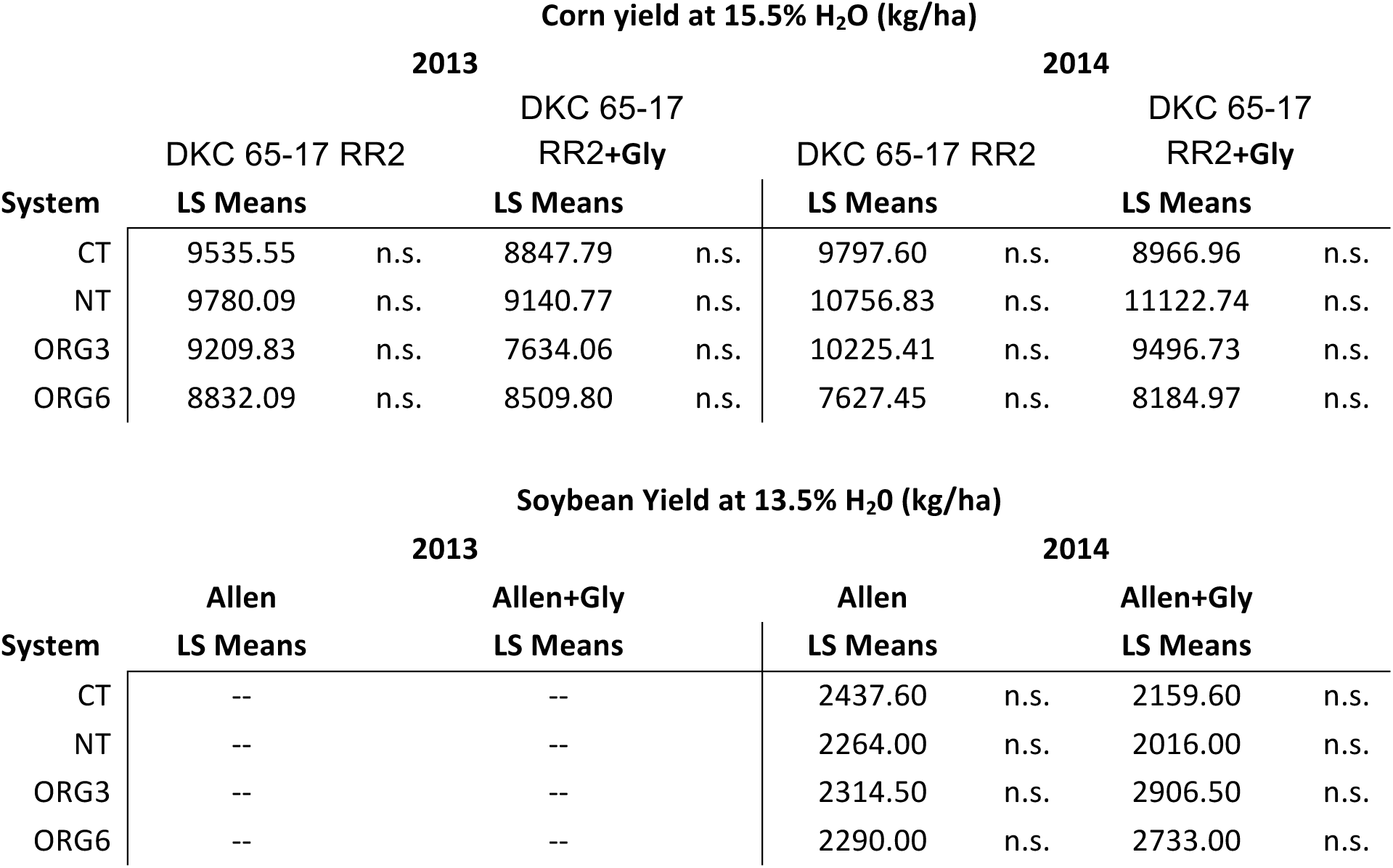
Corn and Soybean yield (kg/ha) for glyphosate treated or untreated plots in chisel till (CT), no-till (NT), Organic 3 yr. rotation (Org3) or Organic 6 yr. rotations (Org6). Comparison of means was calculated within each system for the glyphosate resistant genotype either treated with Glyphosate (Gly) or not. In 2013 an error in microplot harvesting resulted in mixing of treated and untreated plots therefore making the yield data un-usable.

## Discussion

The structure of prokaryote and fungal communities among farming systems and between sampling dates were not driven by glyphosate use. Instead, tillage and carbon inputs appear to be the primary drivers of soil microbiome structure. For instance, even though the Beltsville site had a common history of no-till management prior to 1996, microbial communities today are easily differentiated by farming system. Differences in management have effects that extend beyond microbial taxa to include nematodes [37], as well as soil organic matter and phosphorous concentrations, greenhouse gas emissions, and total energetic costs of the farming system [24, 38, 39].

The absence of glyphosate effects in naïve soil communities suggests that typical application rates of glyphosate do not alter the overall microbial community. Existing literature suggests most microbial communities are susceptible to disturbance, although bias against reporting of no treatment could affect this view [40]. Understanding the factors contributing to resistance of microbes in agroecosystems remains an important goal [41]. In the current study resilience to glyphosate spray could be linked to several factors. Some bacterial and fungal species are known to metabolize glyphosate, and the presence of these organisms may protect susceptible species [42, 43]. Studies reporting effects of glyphosate on soil microbes often use higher concentrations of the herbicide than the approved rate, which may overwhelm buffering by resistant members. Concentration-dependent effects of glyphosate on soil microbial respiration and biomass have been reported and are consistent with reports on other agrochemicals, showing only transient effects at recommended application rates [44].

Greenhouse studies with GR wheat conducted in the Pacific Northwest found only minor effects of glyphosate on microbial communities, and determined site was a major driver of soil microbial community structure [20, 21]. While these studies did detect effects of glyphosate on the prevalence of a few microbial taxa, they applied glyphosate at twice the recommended rate, increasing the likelihood that the microbial community experienced a significant effect. These methodological differences may account for the detection of an effect on the abundance of some taxa after glyphosate exposure where none was detected here, and ultimately increase confidence in our finding that glyphosate has minimal effect on the microbial community when applied at the recommended rate.

The Beltsville and Stoneville sites differ in soil chemistry and physical characteristics (OM, pH). Soil microbial communities in these soils also differ considerably between sites (Figure 1), with Beltsville having higher overall fungal richness and Stoneville having higher prokaryotic richness (Figure 1). The higher richness of fungi and prokaryotes in Beltsville NT plots relative to the other Beltsville management types receiving tillage is consistent with previous studies, and may be due to the spatial heterogeneity that develops over time in the absence of tillage [45]. However, in spite of differences in microbial communities between sites and among management histories, fungal and prokaryotic richness were unaffected by applications of glyphosate in all management and crop treatments.

Community richness changed across the growing season regardless of glyphosate concentration (Figure 4). These results are similar to those of Hart et al. [46] in which the GR corn and its genetically close isoline were grown for one season in Canada and the microbial community compared by TRFLP with and without glyphosate application, although this study could not have tested the long term legacy of glyphosate application. This study also found that seasonality was a significant controlling factor in microbial community structure with and without glyphosate under field conditions.

Previous culture-based work has found that *Fusarium* abundance increased and *Pseudomonas* abundance decreased with glyphosate use [11]. In those studies, *Fusarium* were presumed to be pathogenic while *Pseudomonas* were presumed to be symbionts. However, our metabarcoding and culture data failed to detect an effect of glyphosate on the abundance of any *Fusarium* or *Pseudomonas* spp. And while both barcoding and culture surveys detected other pathogens, none responded to glyphosate (Supplementary Data). Our results are consistent with previous metabarcoding studies [20, 21].

It is important to note that the ITS and 16s gene loci fail to resolve diversity at an adequate level to differentiate pathogenic genotypes from closely related non-pathogenic genotypes [47, 48]. For example, several species of *Metarhizium* known to occur at this site [49] were not represented in the samples from this study. Most likely pathogenic strains were not detected in this study. However, even if they are not identified to the strain level, pathogenic species contribute to the relative abundance of their constituent OTU, and we did not detect any change in total numbers of *Fusarium* spp. OTUs associating with crops due to glyphosate application. This holds true for other genera of pathogenic fungi such as *Alterneria* spp. and *Macrophomina* spp. (supplementary data). It should also be noted that while *Pseudomonas* spp. are often taken to be inherently beneficial, there are at least a few confirmed pathogens [48]; and the type of beneficial function may differ substantially across strains. Regardless, as with fungi, no *Pseudomonas* spp. changed in prevalence as a result of glyphosate treatment.

We also found no reductions in yield by glyphosate application on GR corn or GR soybean in fields with a long history of glyphosate use or with no history of glyphosate use [50, 51]. In a similar study with GR sweet corn, there was even a slight increase in yield associated with glyphosate application [52], which could have been due to hormesis, a phenomenon with non-phytotoxic doses of glyphosate [53]. Lack of effects on yields are consistent with no substantial detrimental effects on rhizosphere microbes.

Although glyphosate is widely used across the globe, relatively few studies have investigated the effect of this herbicide on soil microbial communities in cropping systems with and without a legacy of glyphosate application. This work provides an important contribution into determining the effect of glyphosate on bacterial and fungal communities found in soils. No changes due to glyphosate, coupled with a trend towards higher species richness in no-till plots, suggests this widely employed management practice is not at risk of altering soil microbial communities in a negative manner. However, increased glyphosate application rates in response to evolution of resistant weeds could alter this conclusion. Whether the species richness of no-till systems translates to increases in ecosystem function supportive of crop productivity remains to be fully elucidated.

## Acknowledgments

Sarah Emche assisted with DNA extraction and sequencing preparation, root plating and isolate identification. Other people helped and they are…

## Conflict of interest

The authors declare they have no conflict of interest.

